# Uncovering senescent fibroblast heterogeneity connects DNA damage response to idiopathic pulmonary fibrosis

**DOI:** 10.1101/2025.11.26.690792

**Authors:** Jun-Wei B. Hughes, Akshay Pujari, Anja Sandholm, Duncan Croll, Lea B. Monk, Isaac Joshua, Rachel Butterfield, Cory Horton, Kevin Schneider, Fiona Senchyna, Ian Brown, Ana L. Coelho, Tsung-Che Ho, Hideto Deguchi, Claude Jordan Le Saux, Stefanie Deinhardt-Emmer, Lisa M. Ellerby, Alberto Vitari, David Furman, Cory M. Hogoboam, Alex Laslavic, Pierre-Yves Desprez, Marco Quarta, Judith Campisi

**Affiliations:** Buck Institute for Research on Aging, Novato, CA, USA; University of Southern California, Los Angeles, CA, USA; Rubedo Life Sciences, Inc., Sunnyvale, CA, USA; Cedars-Sinai, Los Angeles, CA, USA; University of California San Francisco, San Francisco, CA, USA; Institute of Medical Microbiology, Jena University Hospital, Jena, Germany; California Pacific Medical Center, Research Institute, San Francisco, California, USA

**Keywords:** Senescence, IPF, DNA damage, heterogeneity, WGCNA

## Abstract

Cellular senescence is a largely heterogeneous state of cell stress that deleteriously accumulates with age. Many types of heterogeneity in senescence have been described; however, cellular senescence within the same cell type has only started to be documented. Here, we show primary, human lung fibroblasts from donors who are healthy or diagnosed with idiopathic pulmonary fibrosis (IPF) exhibit a subtle form of heterogeneity over time after DNA damage. Moreover, senescent IPF lung fibroblasts display a dysregulated transcriptional–protein DNA damage response (DDR). Weighted gene correlation network analysis (WGCNA) reveals unique and known targets linking senescent IPF lung fibroblast heterogeneity to genes associated with DNA damage and repair, cytokine and chemokine responses, and extracellular matrix (ECM) signaling. We combine our healthy and IPF senescent gene expression signatures to develop a novel gene set of senescence-associated genes that identify disease-relevant cells in human single-cell RNA-seq (scRNA-seq) data. Collectively, our results uncover human-relevant senescence signatures, highlight IPF-specific DDR, cytokine and chemokine, and ECM targets, and expand our understanding of how a dysregulated DDR contributes to senescent cell heterogeneity in IPF.

## INTRODUCTION

Cellular senescence is canonically defined by cell cycle arrest, resistance to apoptosis, and the adoption of a senescence-associated secretory phenotype (SASP)^1^. However, exceptions to each of these classical definitions exist as it becomes increasingly evident how heterogeneous cellular senescence is depending on the induction method, cell type, or even microenvironment^2–7^. Here, we describe a novel form of senescence heterogeneity driven by disease state within the same cell type.

IPF is a progressive and fatal lung disease of unknown origin associated with fibrosis and declining lung function^8–10^. Median survival is approximately 3–5 years, with few effective treatments and no cures other than lung transplantation^8–10^. While the etiology of IPF is poorly understood, IPF is largely an age-related disease thought to be driven by persistent injury response and damage repair signaling from alveolar type II (AT2) cells, lung fibroblasts, and to a lesser extent, endothelial cells^11–16^. Both AT2 and fibroblast senescence has been implicated as a driver of chronic matrix deposition and profibrotic SASP signaling that sustains epithelial injury^17–24^. Much research has focused on deep genetic and phenotypic profiling of AT2 cells and lung fibroblasts implicating senescence, dysfunctional wound healing, and dysregulated ECM formation in both of these cell types as drivers of disease progression^25–29^. However, less research has studied how these cells, particularly fibroblasts, respond to stress, which is crucial in an IPF lung niche that is aged and highly inflammatory. In addition, even though past literature has shown primary lung fibroblasts from IPF donors display features of senescence, these cells still divide *in culture* at an equal rate to primary lung fibroblasts from healthy donors, as shown by the Fully-Automated Senescence Test (FAST), (Supp. Fig. 1)^30^. This indicates IPF lung fibroblasts have not “fully” adopted a senescent phenotype and may not completely model a senescent lung fibroblast *in vivo*. An increased understanding of senescent cell heterogeneity in IPF can help build better models for IPF and downstream, improved targets for therapeutic development.

Here, we focus on primary human lung fibroblasts and show largely similar pathway level responses between healthy and IPF lung fibroblasts after senescence induction with more subtle heterogeneity at the individual gene level in senescent IPF lung fibroblasts. We show that after inducing healthy or IPF lung fibroblasts to senesce by DNA damage, our novel gene signature can uniquely identify disease-relevant cell types from human, scRNA-seq data and additionally, is a distinct set of genes relative to other known senescence-associated gene sets. We also determine that IPF lung fibroblasts after DNA damage exhibit lower DNA damage signaling relative to healthy lung fibroblasts at the RNA level but increased damage and repair sites at the protein level. Lastly, we use WGCNA to identify novel and known genes that may drive IPF-specific dysregulation in the DDR and heterogeneity of senescence-associated responses.

## RESULTS

### Healthy and IPF lung fibroblasts display similar pathway level responses to senescence induction

To investigate senescent cell heterogeneity in IPF, we induced lung fibroblasts to senesce from 7 healthy and 9 IPF human donors by γ-irradiation (IR) (Fig. 1A), a common method to induce senescence primarily through DNA double-strand breaks (DSBs)^5,17,31–33^. All analyses were performed relative to donor-matched non-irradiated controls to isolate senescence-specific responses. Bulk RNA-sequencing (RNA-seq), the Assay for Transposase-Accessible Chromatin using sequencing (ATAC-seq), and FAST were performed 10-days after IR^30^. FAST shows that healthy and IPF lung fibroblasts both divide at similar rates and neither are positive for senescence-associated beta-galactosidase (SA-β-gal) staining or negative for 5-ethynyl-2′-deoxyuridine (EdU) incorporation prior to IR (Supp. Fig. 1). While previous scRNA-seq studies show IPF lung fibroblasts exhibit some features of senescence, FAST uncovers that other features of senescence, such as cell cycle arrest, are missing without senescence induction, which is shown in Supp. Fig. 1 by the number of cell cycle arrested lung fibroblasts increasing after senescence induction in both healthy and IPF conditions^25,26,28^. In addition, Supp. Fig. 1 also shows that “senescent-like” cells (classified by EdU- and SA-β-gal+ staining) in healthy or IPF lung fibroblasts are not present in the RNA-seq data prior to IR, suggesting that senescence-associated signatures present in the healthy and IPF lung fibroblasts after IR are indeed from senescence induction. Differential expression analysis shows hundreds of differentially expressed genes (DEGs) between healthy and IPF conditions after IR relative to the mock-irradiated control (CTL) cells (Fig. 1B). Principal component analysis (PCA) shows biological heterogeneity between donors drive separation between data points as well as due to IR treatment (Fig. 1C). While there is no distinct separation between healthy and IPF disease conditions, Fig. 1D shows 910 upregulated and 469 downregulated DEGs uniquely in IPF lung fibroblasts after IR. Fig. 1E however, shows that the majority of genes up and downregulated after IR in both healthy and IPF lung fibroblasts are similar, consisting of pathways relevant to senescence induction such as p53 transcriptional gene network upregulation and cell cycle and DNA damage response-associated pathway downregulation. Our data shows that senescence induction in healthy and IPF lung fibroblasts drives largely similar pathway level responses with more subtle heterogeneity at the individual gene level.

**Figure 1.**
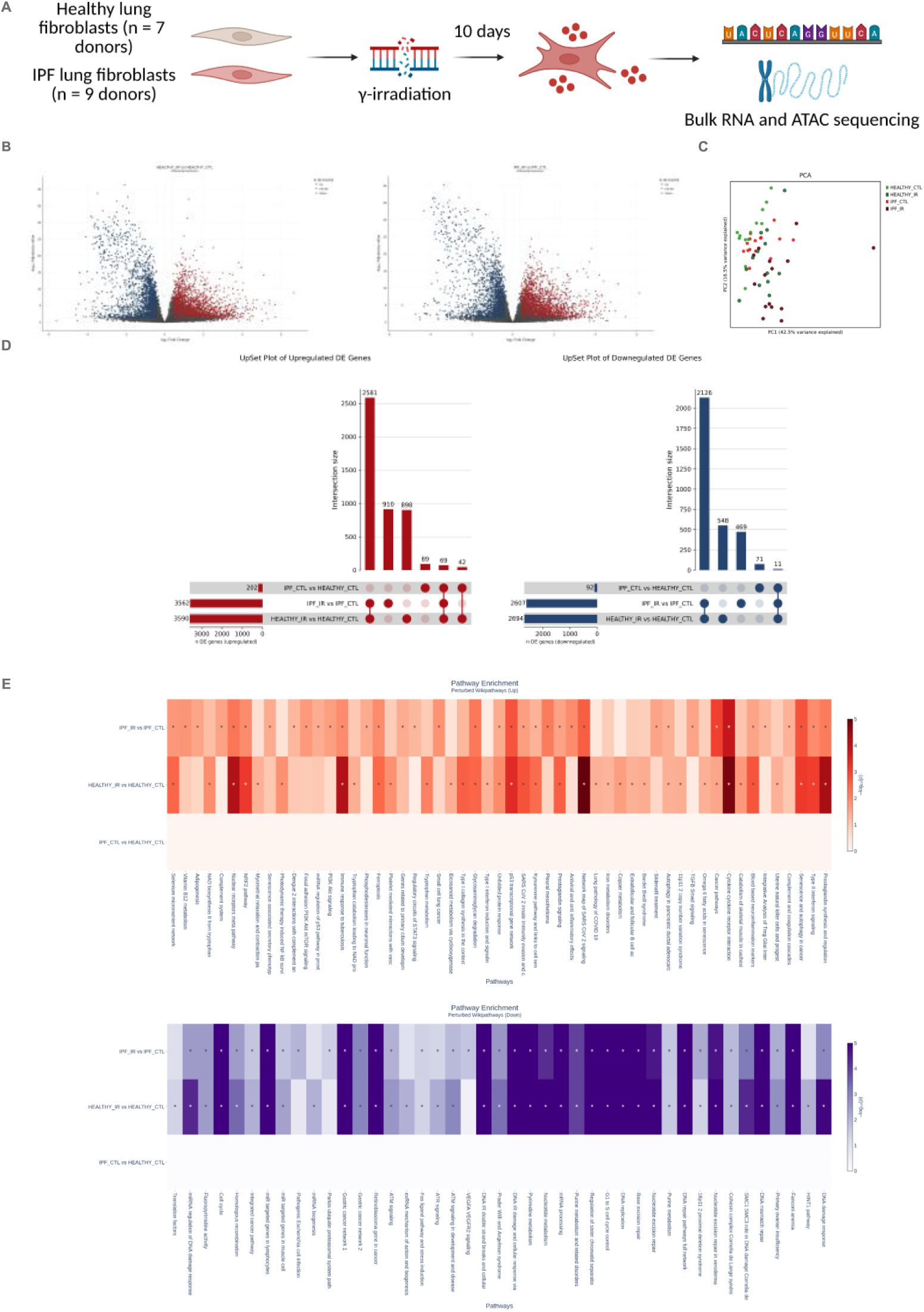
Healthy and IPF lung fibroblasts activate similar pathways in response to senescence induction. (A) Schematic of 7 healthy and 9 idiopathic pulmonary fibrosis (IPF) primary, human, lung fibroblasts, between ages 29-75, induced to senesce by γ-irradiation (IR). RNA and DNA were collected for bulk RNA sequencing (RNA-seq) and ATAC-seq, respectively, 10-days post IR. (B) Volcano plot of differentially expressed genes (DEGs) in healthy IR vs CTL lung fibroblasts (left) and IPF IR vs CTL lung fibroblasts (right). DEG cutoffs were defined by log2FC < -0.4 or > 0.4, p-adjusted value < 0.05, and minimum expression of 4 transcripts per million. (C) Principal component analysis (PCA) of healthy and IPF lung fibroblasts with and without IR. (D) UpSet plots displaying the number of unique and overlapping genes between healthy and IPF lung fibroblasts with and without IR. (E) Gene set enrichment analysis performed using Wikipathway gene sets of upregulated (top) and downregulated (bottom) DEGs in healthy and IPF lung fibroblasts after IR.

### DNA damage response signaling is dampened in IPF over time after IR

Given the similarities between healthy and IPF lung fibroblasts at a single time point after senescence induction, we asked if IPF lung fibroblasts may respond differently than healthy lung fibroblasts over time after IR. We profiled early time points reasoning that early responses to DNA damage may drive later cell state changes^6,34,35^. We performed RNA-seq on 6 healthy and 6 IPF donor lung fibroblasts at 0, 8, 24, and 48 hours and 10-days (240h) after IR (Fig. 2A-B). As expected at the 8-hour timepoint, both healthy IR and IPF IR lung fibroblasts, relative to their respective controls, increased DDR signaling and decreased canonical pathways associated with cell cycle and mitosis (Fig. 2C). However, when comparing IPF IR to healthy IR 8-hours after IR, we observed a lower enrichment of pathways associated with DNA damage, such as the E2F target pathway which is a transcription factor (TF) family that targets many cell cycle and DNA repair associated genes (Fig. 2C). It is important to emphasize that IPF lung fibroblasts still upregulated pathways associated with DDR and DNA repair 8-hours after IR; however, this upregulation was to a lesser extent than healthy lung fibroblasts 8-hours after IR. Investigating which E2F target genes were downregulated in IPF lung fibroblasts showed many genes involved in DDR signaling and DNA repair, such as PCNA, RPA3, and SHMT1, which all play a role in repairing DSBs (Fig. 2D)^36^. To look at how E2F target genes and DNA repair associated genes contribute to the overall transcript pool over time after IR, we investigated relative transcript abundances between healthy and IPF lung fibroblasts over time after IR (Fig. 2E). We show that crucial gene transcripts involved in DDR and DNA repair, such as PCNA and ERCC2, do not increase in abundance 8-hours after IR in IPF lung fibroblasts relative to healthy lung fibroblasts (Fig. 2E)^37^. Ultimately, IPF lung fibroblasts show a decreased DDR relative to healthy lung fibroblasts over time after DNA damage at the transcriptional level.

**Figure 2.**
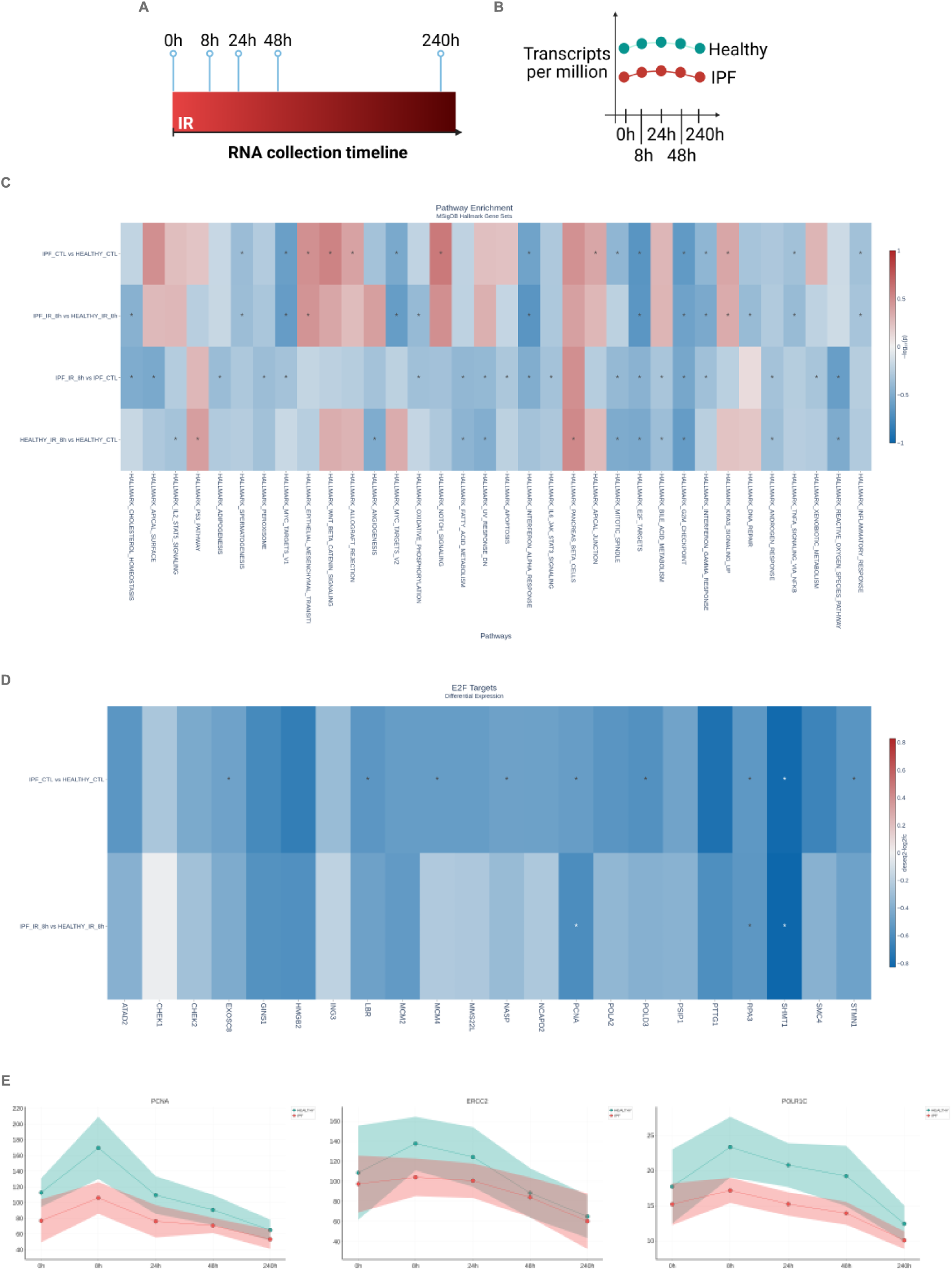
RNA sequencing over time after IR in IPF lung fibroblasts reveals lower transcriptional DNA damage response. (A) Schematic of RNA collection time points at 0, 8, 24, 48, and 240-hours after IR. 6 healthy (donors: AJAZ, HLF250, AJDL, HLF872, HLF763, AJIQ) and 6 IPF (donors: 83A, 110A, IPF809, 86A, 127A, 155B) lung fibroblasts were used for this time course analysis. (B) Schematic for (E) detailing transcript counts at 5 time points prior to and after IR in healthy (turquoise line) and IPF (red line) lung fibroblasts. (C) Gene set enrichment analysis performed using The Molecular Signatures Database (MSigDB) hallmark gene set collection with DEGs from healthy and IPF lung fibroblasts at timepoints 0h prior to and 8h after IR. (D) E2F gene target differential expression heatmap with DEGs from healthy and IPF lung fibroblasts at timepoints 0h prior to and 8h after IR. (E) Relative transcript abundance of genes associated with DNA repair over time after IR between healthy and IPF lung fibroblasts.

### DNA damage response and repair signaling at the protein level is increased in IPF lung fibroblasts over time after IR

To confirm decreased DDR and DNA repair signaling at the mRNA level was similar at the protein level, we performed immunocytochemistry (ICC) over time after IR with 3 healthy and 3 IPF donor lung fibroblasts (Fig. 3A). We chose earlier time points than in the RNA-seq time course paradigm since most protein response to DNA damage occurs immediately, at the post-translational level, and with proteins already translated to repair damaged DNA as quickly and efficiently as possible^38^. We stained for proteins that localize to DNA damage sites, such as γH2AX and phosphorylated ATM (pATM), and proteins that repair DNA damage, such as RAD51 and PCNA (Fig. 3C/D)^38^. We imaged and analyzed these markers automatically using the Harmony’s Operetta CLS system and High-Content Analysis software, respectively. In brief, Harmony’s analysis software uses DAPI staining to segment the nuclei in each image and then nuclear intensity and foci number were quantified based on nuclear segmentation and algorithmic detection of nuclear foci (Fig. 3B). As expected, γH2AX and pATM intensity and foci levels were increased 15 minutes after IR in both healthy and IPF lung fibroblasts (Fig. 3E-F).

**Figure 3.**
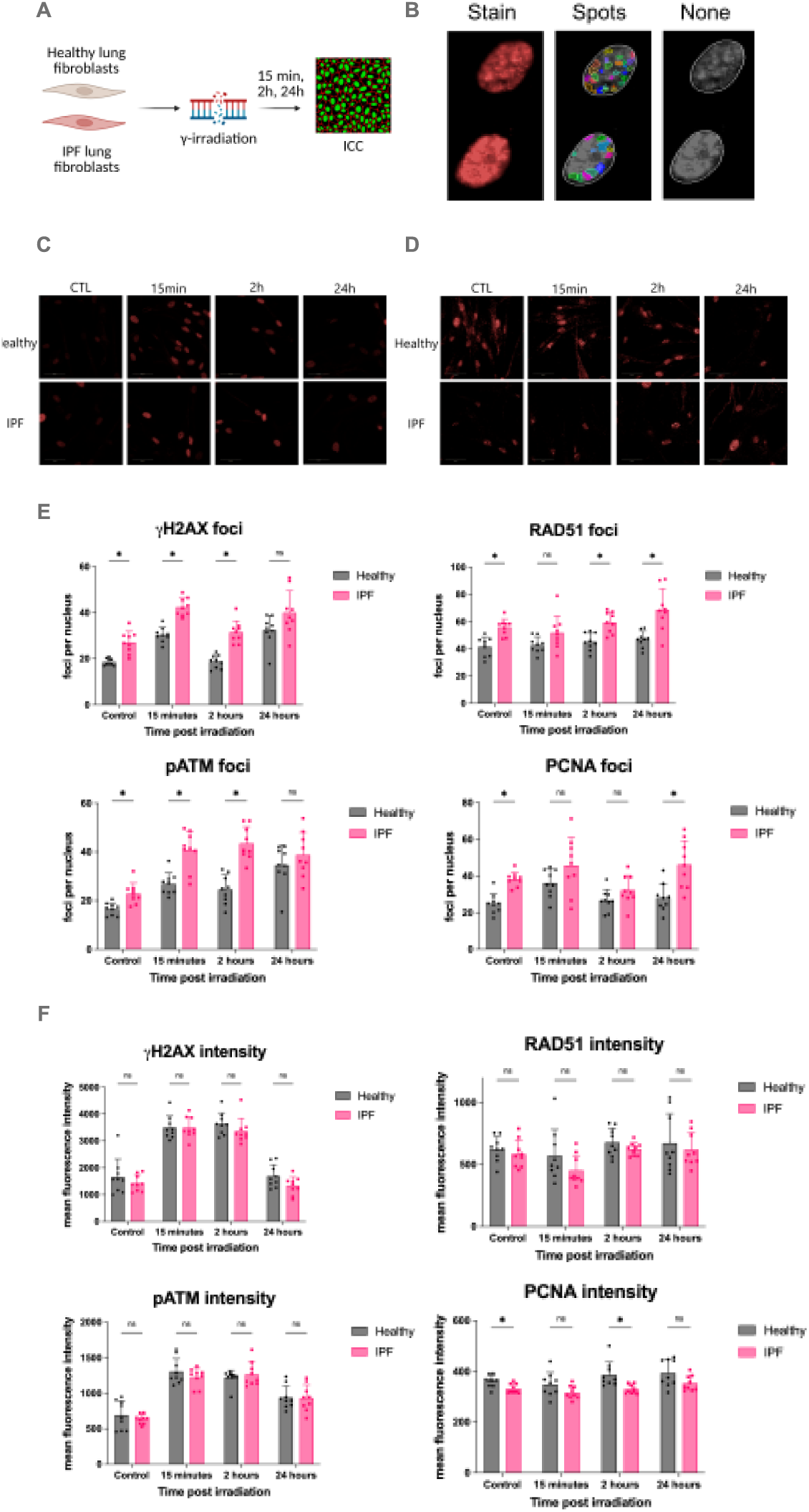
IPF lung fibroblasts display higher damage accumulation and DNA repair protein recruitment prior to and over time after IR. (A) Schematic of immunocytochemistry (ICC) performed after IR in healthy and IPF lung fibroblasts. 3 healthy (donors: AJAZ, HLF250, AJDL) and 3 IPF (donors: 83A, 110A, 112A) lung fibroblasts were used for the time course analysis. (B) Harmony spot software analysis example of γH2AX foci in two nuclei at 63X magnification. Staining is in the left panel is a red γH2AX stain and the “Spots” middle panel is the software identifying foci in the nucleus from the γH2AX staining. Grayscale image of the red stain provided in the right panel showing algorithmic nuclear segmentation. (C-D) Healthy and IPF lung fibroblasts were fixed and stained by ICC for γH2AX (C), Rad51 (D), pATM, and PCNA at timepoints 15 min, 2h, and 24h after IR (pATM and PCNA images shown in Supp. Fig. 9). Images taken on the Harmony Operetta CLS system at 63X. (E-F) Foci and nuclear intensity quantification of images performed by the Harmony analysis system and Harmony spot finder analysis, respectively. Statistics were calculated using an unpaired t-test with Welch correction on each row and Holm-Šídák’s multiple comparisons test with alpha = 0.05.

Surprisingly, in IPF lung fibroblasts prior to and over time after IR, both DNA damage sites, quantified by γH2AX and pATM foci, and repair proteins, quantified by RAD51 and PCNA foci, were increased relative to healthy lung fibroblasts (Fig. 3E). Nuclear intensity analysis of γH2AX, pATM, RAD51, and PCNA to infer protein levels by ICC did not show any differences between healthy and IPF lung fibroblasts (Fig. 3F). Although IR does not largely result in single nucleotide polymorphisms (SNPs), we looked into whether a differential burden of SNPs in IPF lung fibroblasts prior to IR may explain the dysregulated DDR^39–41^. However, our SNP analysis showed no differences between healthy and IPF lung fibroblasts except at 8-hours after IR in healthy lung fibroblasts (Supp. Fig. 2). In contrast to the mRNA level response to DNA damage, proteins recruited to damage sites and responsible for DNA repair after DNA damage are upregulated in IPF lung fibroblasts suggesting a dysregulated transcriptional and protein level response to DNA damage in IPF lung fibroblasts.

### Healthy and IPF senescent lung fibroblast gene set identifies unique disease-relevant cell populations within single-cell IPF data

To understand the clinical relevance of our *in culture* model, we compared our healthy and IPF DEGs over time after IR to human scRNA-seq data from Habermann et. al. 2020 which sequenced 10 healthy and 10 IPF donor lungs^25^. We used a scoring algorithm which sums the expression z-scores between our RNA-seq data and the scRNA-seq data (Fig. 4A). Cell types from the scRNA-seq data that are most similar in gene expression to our *in culture* gene expression would have the highest summed z-scores (Fig. 4A). We compared the scRNA-seq data to both our healthy IR and IPF IR gene expression data since the pathway level responses were largely similar, which was further emphasized by that each condition picked up similar populations in the scRNA-seq data (Supp. Fig. 3A-B). The IR gene signature that best mapped to disease-relevant single-cell clusters was determined to be a subset of 163 genes from our entire time course of healthy IR and IPF IR DEGs. These 163 genes preferentially map to senescent-like clusters from the scRNA-seq data and partially are composed of known senescence-associated genes, such as CDKN1A and CCL2 (Fig. 4B-C). Notably, the gene signature most resembles cluster 35 in the early response to IR and then more resembles cluster 14 at the later time points after IR (Fig. 4C). Cluster 14, which is composed of PLIN2+ mesenchymal cells and myofibroblasts from interstitial lung disease, was previously identified in Habermann et. al. 2020 as potential populations of interest in the human IPF lung for future therapeutic development (Fig. 4C)^25^. We build on their study by uncovering more senescence-associated markers used to locate the disease-associated PLIN2+ cell populations in human scRNA-seq data (Fig. 4C). Cluster 35 seems to be a novel cluster of mesothelial cells identified by the IR gene signature that is highly expressing both CDKN1A and CDKN2A. To better understand the genes that make up the IR gene signature, we ran SASP pathway enrichment on the healthy IR and IPF IR gene sets 10-days after IR. Many of these SASP genes were upregulated in the healthy IR and IPF IR gene sets; however, when comparing the 163 gene disease-relevant IR gene signature to the SASP and SenMayo gene sets, a gene set known to identify senescent cells across tissues, we found little overlap between each of the gene sets^42^. This emphasizes the need for further gene sets to better identify senescent cells across human tissues and shows we can identify disease-relevant cell populations in the human lung using our novel gene set.

**Figure 4.**
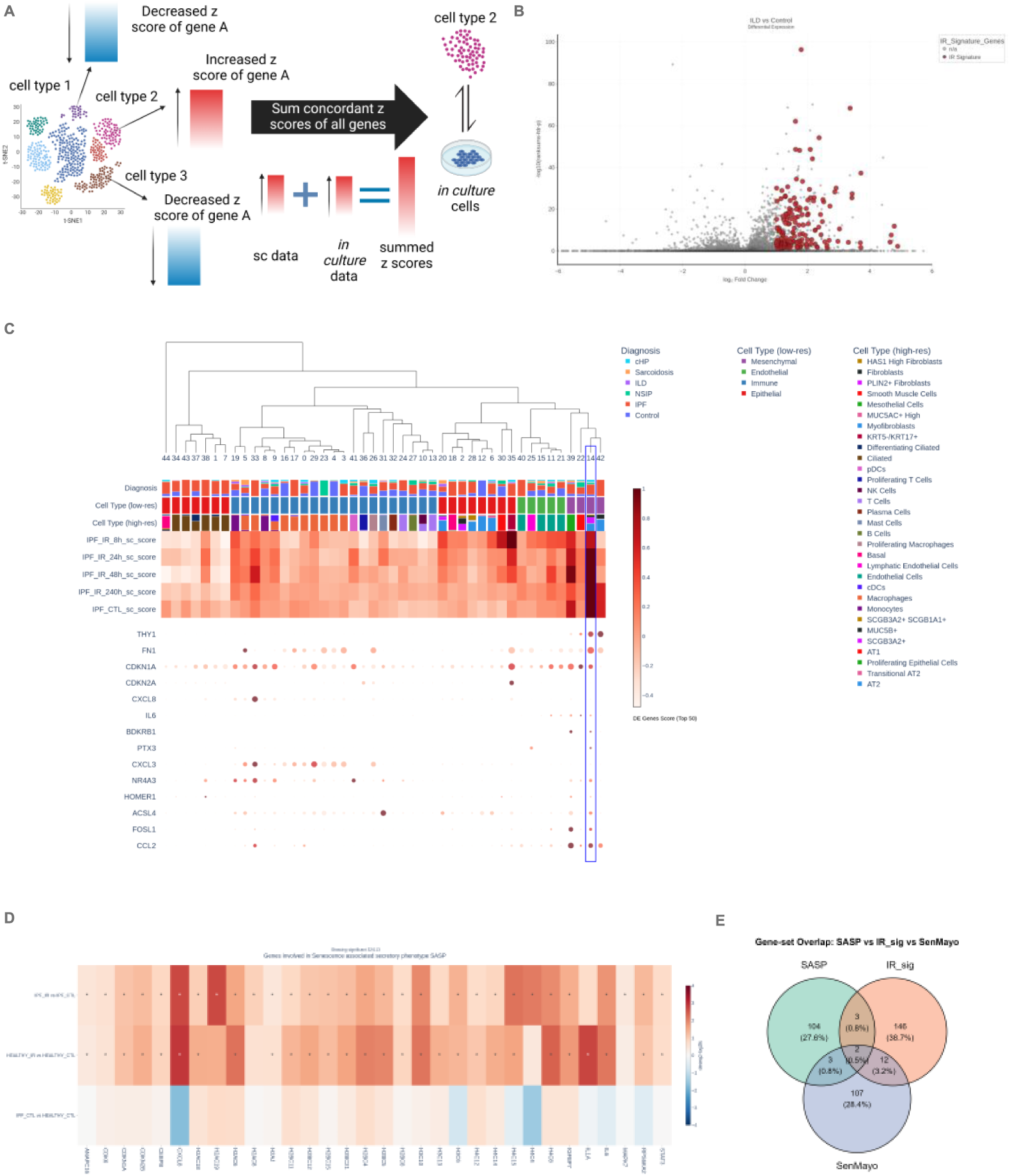
Scoring algorithms reveal a disease-relevant gene signature from healthy and IPF senescent cell data within human scRNA-seq data. (A) Diagram of z-score comparisons between RNA-seq and scRNA-seq data. In brief, RNA-seq data has gene expression values converted to z-scores which is then summed against the z-scores from each cell type gene expression in the scRNA-seq data. The cell type from the scRNA-seq data with the highest summed z-score is the most similar, in terms of gene expression, to the *in culture* data. (B) Volcano plot of healthy and IPF RNA-seq data after senescence induction. Red bubbles are genes are listed in (C) used to identify cell populations in the scRNA-seq data comprising of the top 50 differentially expressed genes in our senescent healthy or IPF lung fibroblast data sets. (C) Hierarchical clustering of scRNA-seq data from Habermann et. al. 2020. The first row of columns indicates diagnosis (cHP: Chronic hypersensitivity pneumonitis, Unclassifiable ILD: interstitial lung disease, IPF: Idiopathic Pulmonary Fibrosis, NSIP: Nonspecific Interstitial Pneumonia). The data is then clustered based on cell type (low-res) from the scRNA-seq data in the second row of columns. The third row of columns indicates further cell type (high-res) detail based on gene expression markers. The darker the red color indicates more similarity to the scRNA-seq cluster. Outlined column (number 14) indicates a potential column of interest due to high similarity scores. (D) Wikipathway enrichment of senescence-associated secretory phenotype (SASP) related genes for all healthy and IPF DEGs. (E) Venn diagram showing the overlap between the SenMayo, Wikipathay SASP, and the healthy and IPF IR (IR_sig) gene sets. The IR gene signature is comprised of 163 total genes which is a subset of the entire healthy IR and IPF IR DEG list that showed the most consensus signature to the scRNA-seq ILD gene expression signature.

### WGCNA reveals IPF lung fibroblast senescence is associated with dampened DNA repair and cytokine signaling but increased extracellular matrix signaling

Although healthy and IPF lung fibroblasts after senescence induction seem largely similar, IPF lung fibroblast heterogeneity over time after DNA damage and dysregulation between mRNA and protein level responses to DNA damage warranted further exploration of genes that may be driving these differences. We used WGCNA combined with our differential expression analysis to determine which genes may drive the subtle heterogeneity in the IPF IR phenotype^43^.

Importantly, our sample number gave us confidence that the WGCNA would reach sufficient power for meaningful results, as Langfelder and Horvath recommend using at 15 samples and our total sample number was 16 with timepoints and two biological replicates per sample^43^. It should also be noted that meaningful results can still be discovered with less than 15 samples, which emphasizes the importance of validating WGCNA module genes regardless of sample size^44^. In brief, WGCNA uses transcript counts to build a gene co-expression network which is then used to determine which genes are highly correlated with a specific condition, which in our case was IPF over time after IR^43^. The WGCNA then outputs gene modules and how they are related to each other, in addition to a heatmap with z-scores of how correlated each gene module is to a specific condition (Fig. 5A)^43^.

**Figure 5.**
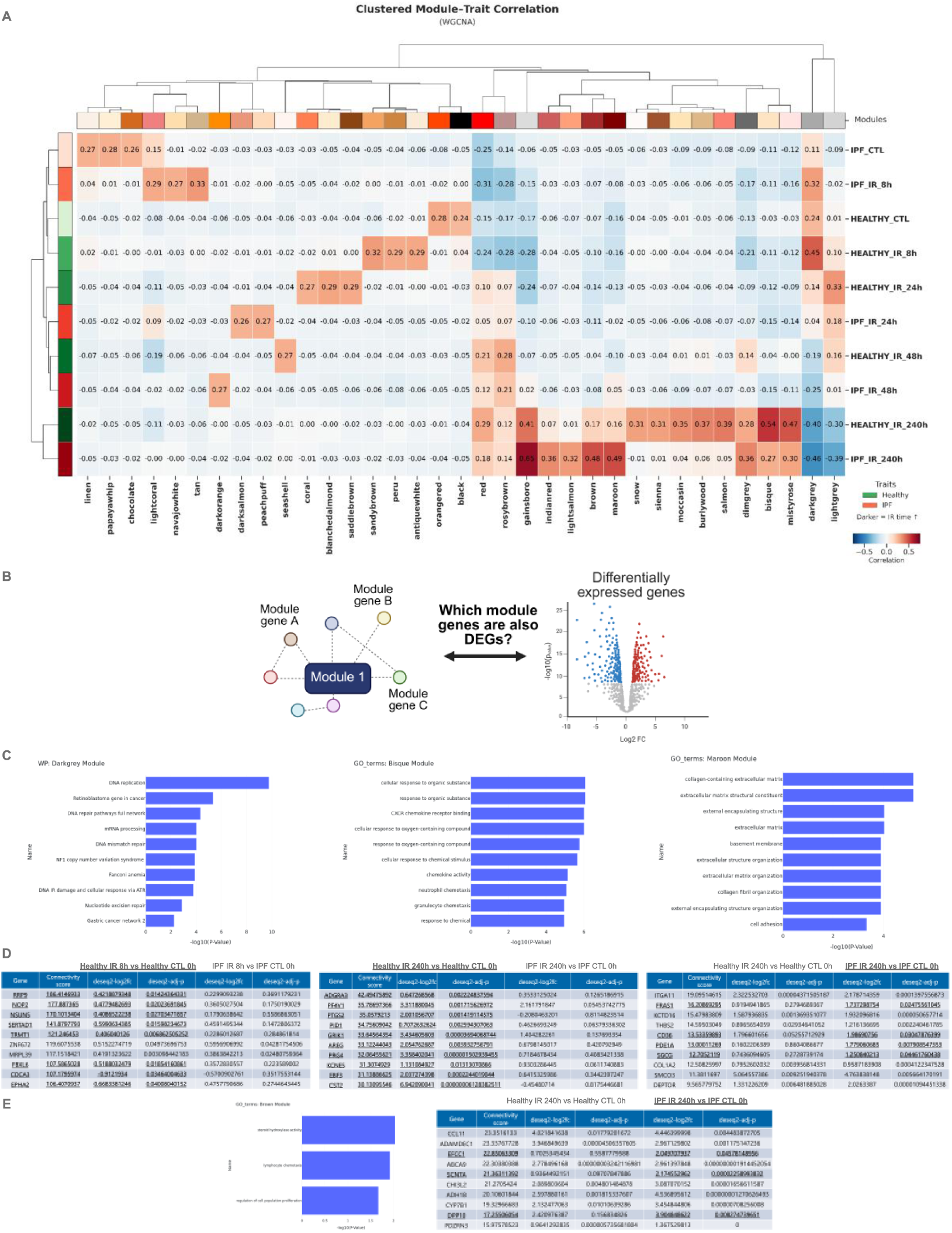
Weighted gene correlation network analysis (WGCNA) highlights gene modules related to DNA repair, cytokine signaling, and the extracellular matrix that are correlated to IPF lung fibroblasts after IR. (A) WGCNA module-trait heatmap relationships. Each row is a trait and each column is a module eigengene. Each cell includes corresponding correlation to the specific module and correlation is indicated by the value in the box. Correlation ranges from -1 to 1 and blue indicates a negative correlation while red indicates a positive correlation. (B) Schematic detailing hub gene identification by comparing module genes to DEGs to determine which genes overlap. (C-E) Wikipathway or GO term enrichment of genes from the Darkgrey, Bisque, Maroon, and Brown modules with accompanying, overlapping module genes and DEGs given differential correlation between healthy and IPF at 8h or 240h after IR. Top 10 partially overlapping module genes and DEGs shown in the adjacent tables. DEGs present in the healthy or IPF senescent conditions, relative to their respective controls, are bolded and underlined as potential hub genes correlating to the healthy or IPF phenotype. Connectivity scores determined by the pyWGCNA package and log2fc and adj-p values determined by DESeq2.

Our WGCNA shows many highly and differentially correlated modules with healthy or IPF conditions over time after IR (Fig. 5A). To add biological relevance to the results from the WGCNA, we compared the gene module lists to our differential expression gene lists, outputting only module genes that were also significantly differentially expressed (Fig 5B). In addition, Wikipathway and GO term analysis was run on each gene module list from the WGCNA (Fig. 5C-E). We highlight 4 gene modules from the WGCNA that were differentially correlated between healthy and IPF conditions (Fig. 5C-E). Of immediate interest was the Darkgrey module, which showed enrichment in DNA damage and repair pathways and was correlated more to healthy lung fibroblasts 8-hours after IR than IPF lung fibroblasts (Fig. 5C-D). Many of the highly correlated genes to the healthy 8-hours after IR lung fibroblast condition were also DEGs but not DEGs in the IPF condition (Fig. 5C-D). This data is similar to earlier data suggesting a transcriptionally dampened DDR in IPF lung fibroblasts after DNA damage. The Bisque module was also of interest as the module was highly correlated with healthy lung fibroblasts, and not IPF lung fibroblasts, 10-days after IR, even though the GO-enriched pathways were mainly related to cytokine and chemokine signaling (Fig. 5C-D). Some of the overlapping Bisque module genes and DEGs shown in Fig. 5D are known to be upregulated in the IPF lung, such as PTGS2 and AREG^45–47^. In contrast to the Darkgrey and Bisque modules, the Maroon and Brown modules were correlated with IPF lung fibroblasts 10-days after IR and showed enrichment in ECM-related pathways and steroid hydroxylase activity, respectively (Fig. 5C-E). While ECM remodeling in the IPF lung is known to be highly relevant to disease progression, many of our overlapping module genes and IPF specific DEGs in the Maroon and Brown modules have not been directly linked to IPF yet, such as FRAS1, SGCG, and DPP10 (Fig. 5C-E). In summary, the WGCNA identifies additional evidence of dampened DDR signaling in IPF at the mRNA level, along with decreased cytokine signaling and increased ECM signaling. Many of the newly identified overlapping module genes and DEGs can be used to validate and potentially modulate IPF-specific phenotypes in the future.

## DISCUSSION

With age-associated IPF rising in prevalence with the increase in our aging population and the disease etiology still unknown, better understanding the potential drivers of IPF, such as the heterogeneity of senescent cell responses in IPF, is crucial to developing better targets for the disease. While much work has been done to profile IPF and senescence-associated responses in the IPF lung niche, there remains gaps in the senescence models, such as how IPF lung fibroblasts still divide *in culture* and may not be “fully” senescent. In addition, while stress responses in epithelial AT2 cells have been studied extensively, stress responses in the fibroblast niche, which supports alveolar cell function, have been less studied. Our findings identify a previously underappreciated form of senescence heterogeneity—driven by disease state within a single cell type, the lung fibroblast. In particular, we expose healthy and IPF lung fibroblasts to γ-irradiation and profile DNA damage-induced senescence over time after IR using RNA-seq, ATAC-seq, and FAST. We show mainly similar senescence-associated responses between healthy and IPF lung fibroblasts with more subtle heterogeneity when investigating temporal dynamics of mRNA and protein levels after DNA damage. IPF lung fibroblasts display a dysregulated DDR, potentially resulting from differences between gene and protein level responses to DNA damage. We highlight how we have discovered a novel disease-relevant gene signature that is able to identify unique and known senescence-associated cell clusters in human scRNA-seq data. Finally, we use WGCNA to uncover known and novel IPF-specific targets associated with dampened DDR and cytokine and chemokine signaling and increased ECM signaling.

It could be reasoned that the differences in mRNA and protein levels in response to DNA damage is due to a higher basal state of stress in IPF lung fibroblasts that results in an attenuated transcriptional DDR with heightened damage and repair protein sites; however, this needs deeper mechanistic support^38^. Phospho-proteomics and replication stress markers over the same time points as our RNA-seq time course could provide better insight into potential dysregulation between mRNA and protein level responses in IPF^38^. Previous reports detail genetic overexpression of DNA repair proteins RAD51 or OGG1 in mouse IPF models showed improved IPF phenotypes, such as decreased markers of senescence and fibrosis, so this could be another avenue to determine if relief of damage and repair protein sites results in an overall improvement in IPF-specific phenotypes^31,48,49^.

To attempt to relieve damage sites from IPF lung fibroblasts after DNA damage, we treated IPF fibroblasts with MDM2 inhibitors RG7388 and HDM201 which was previously reported to decrease γH2AX and cytoplasmic chromatin fragments and increase DNA repair^31^. We did observe a decrease in γH2AX intensity and pATM foci after treatment in DNA-damaged IPF lung fibroblasts; however RAD51 intensity and overall cell numbers decreased after treatment in both the CTL and IR conditions, indicating that the MDM2 inhibitors may be pushing the healthy and IPF lung fibroblasts to apoptosis instead of activating DNA repair and relieving damage sites. Follow up experiments should profile p53 levels in healthy and IPF lung fibroblasts to determine if p53 activation is necessary to activate DNA repair as well as assess DNA repair activity and the number of apoptotic cells after MDM2 inhibitor treatment.

Due to technical limitations of ATAC-seq on irradiated lung fibroblasts, our data from healthy and IPF lung fibroblasts after IR were not sufficiently powered. We found that across biological donors, senescence induction by IR halved the number of alignable reads after ATAC-seq, which decreased the power of downstream analyses like pathway and motif enrichment. Our data suggests potential dysregulation in enhancer regions related to DNA damage; however, conclusive results should be followed up by validation of locus-specific accessibility. p53 Chromatin Immunoprecipitation Sequencing (ChIP) might also be a relevant follow up to the ATAC-seq data since p53 is a known protein to alter the DDR chromatin accessibility landscape^50^.

Another limitation to this study is that our “fully” senescent phenotype is not fully developed as we have only assessed EdU, SA-β-gal, and senescence-associated gene expression markers after senescence induction. Future studies should more extensively profile senescence-associated markers, such as p16, p21, Ki67, LMNB1, SASP (IL6, IL8, MMPs, CXCLs), and apoptosis-resistance markers, 10-days after DNA damage as well as 14 to 21 days after DNA damage to establish if the “fully” senescent phenotype is stable. We did test if healthy or IPF lung fibroblasts 10-days after IR had differential susceptibility to ABT263, which targets the anti-apoptotic BCL protein family, and our data suggests increased susceptibility in younger and IPF lung fibroblasts to ABT263 relative to adult lung fibroblasts^51^. However, more extensive follow up should be performed with an increased number of donors and senolytic agents, such as ABT737 and FOXO4-DRI, in addition to a variety of apoptosis assays^52,53^.

The novel IR gene signature should be further validated on more patient scRNA-seq data to better benchmark the gene set against previously deposited disease-relevant and senescence-associated gene sets. We do not rule out that the SASP and SenMayo gene sets are more representative of a different part of our entire healthy and IPF senescence signature; rather, we show that a subset of the entire senescence signature uncovers a consensus IR gene signature that maps to disease-relevant populations in the scRNA-seq data that the SASP and SenMayo gene sets are not able to map to. In addition, it remains to be tested if the IR gene signature is able to stratify slow versus fast progressing IPF patient populations as well as if our more subtle healthy versus IPF gene signature heterogeneity may better identify stratified patient populations. This should then be linked to our WGCNA data to determine if IPF-specific phenotypes, such as dampened DDR and increased ECM signaling, map to patient-specific cell populations.

Although subtle, our WGCNA results reveal a number of known and novel targets that warrant further investigation and validation. Tying changes in DDR to changes in ECM function in IPF lung fibroblasts would provide more mechanistic insight as to how the WGCNA modules play a role in the overall IPF cell state. Notably, many of the DDR genes that are upregulated in healthy lung fibroblasts 8-hours after IR but not IPF lung fibroblasts are also related to ribosomal signaling and function, which could be a novel angle of investigation into how boosting ribosomal function could impact DDR and the ECM in IPF. In addition, DPP10 was identified as an IPF-specific module gene and DEG, which could be of interest for our IPF-specific phenotype characterization based on recent work implicating DPP4 in IPF and senescence^54^. Also of importance is that two overlapping module genes and DEGs, NOP2 and PTGS2, were also in the IR gene signature and not in the SASP or SenMayo gene sets, suggesting drivers of IPF-specific phenotypes present both *in culture* and in human data.

In summary, we have better documented senescent cell heterogeneity in IPF to reveal unique human-relevant targets linked to a dysregulated DDR, dampened cytokine and chemokine signaling, and increased ECM signaling and developed a novel disease-relevant, senescence-associated gene signature that maps to patient scRNA-seq data.

## MATERIALS AND METHODS

### Cell culture

Primary human lung fibroblasts were purchased from Lonza (CC-2512) or Lifeline Cell Technology (FC-0049). Primary human lung fibroblasts were also received from collaborators at the University of California San Francisco or Cedars-Sinai and isolated at their originating institutions through methods previously described^55^. A complete list of details for each donor lung fibroblasts can be found in the “Supplementary Data Donor Information” table. A schematic of the experimental design with each of the donors listed can be found in Supp. Fig. 7. IMR90 fibroblasts were purchased from Coriell (Coriell, I90-10) at low population doubling (PD). Lung fibroblasts were cultured in DMEM (Corning, 01-017-CV) supplemented with 10% FBS (R&D Systems, S11550H), 100 units/mL penicillin and 100 μg/mL streptomycin (R&D Systems, B21210), 10 mM HEPES (Life technologies, 15630080), and 1X GlutaMAX™ supplement (Life technologies, 35050061) at 37°C, 14% O2, 5% CO2. All experiments were performed with the same lot of FBS in full serum. Taking into account that atmospheric oxygen levels (about 20% O2) are different from physiological oxygen levels (3% O2), we chose a 14% O2 concentration for our primary lung cell culture since alveolar oxygen levels are around 14% O2 and primary lung fibroblasts support this niche of epithelial cells^56–58^. Primary lung fibroblasts were cultured in T25 flasks or 6-well or 12-well plates (Genesee Scientific, 25-100) for sequencing experiments or 96-well microplates for microscopy imaging, with media changes every 2–3 days. All cultured cells were monitored and tested regularly for mycoplasma (Lonza, LT07-318). All cultured cells were counted at each split and PDs were monitored over the course of the culture. PDs were kept below 20 for all experiments with no noticeable effects on proliferation before or after PD 20 for all primary cells. PDs were matched between healthy and IPF samples when possible. Healthy (ages 29-73) and IPF (ages 33-75) donors were age-matched between ranges listed as best as possible. Donor samples were not sex-matched and some age and sex information is missing as not listed by collaborators.

### Senescence induction and drug testing

Senescence was induced as by ionizing radiation (IR). For IR-induced senescence, the cells were irradiated at 25-50% confluency with 15 Greys (Gy) and medium change was performed immediately after treatment. Cells were considered senescent after 10-days post IR, during which medium was regularly changed (every 2–3 days). ABT263 (Selleck, S1001) treatment was performed 10-days post IR for 24-hours and cells fixed for FAST analysis after ABT263 incubation. Serial titrations were performed for the five (0.111, 0.333, 1, 3, 9 µM) ABT263 concentrations tested with the DMSO control (Supp. Fig. 6). RG7388 (MedChemExpress, HY-15676) and HDM201 (MedChemExpress, HY-18658) treatment was performed 5 days prior to IR at 100nM and 25nM, respectively, at each media change as previously described (Supp. Fig. 4)^31^.

### Bulk RNA-seq processing

Bulk RNA-seq libraries were generated from total RNA isolated from samples collected at 0h (pre-irradiation) and 8h, 24h, 48h, and 240h post IR. The cells were lysed in the lysis buffer plus 2-Mercaptoethanol (Sigma, M3148-25ML) according to manufacturer’s instructions (VWR, BIO-52073). RNA was isolated according to the protocol and sent out to Novogene for sample QC, library preparation, and sequencing. Sequencing was performed at 20M paired-end reads per sample on the NovaSeq X Plus Series (PE150) (Illumina) through Novogene. FASTQ files provided by the vendor were used as the starting point for computational analyses.

Raw reads were quantified using kallisto, performing pseudo-alignment against a standard human reference transcriptome supplied by the kallisto|bustools resource for Homo sapiens (GRCh38 genome)^59^. Transcript-level count estimates (estimated counts and TPMs) per sample were then summarized to gene-level count matrices using the corresponding transcript-to-gene annotation. The gene-level estimated count matrices along with sample metadata were used for bulk RNA-seq dataset generation and analyses.

### RNA-seq mutation analysis

RNA-sequence mutations (SNPs) were identified using rnaseqmut (https://github.com/davidliwei/rnaseqmut) using sorted BAM files (rnaseqmut {BAM file} followed by rnaseqmut -l ALLMUTLIST.txt {sorted BAM} using the ALLMUTLIST.txt created from all identified SNPs from all files. Files were merged with merge2ndvcf. SNPs were aggregated for each sample that had identified SNPs for each timepoint, disease factor, and sample ID. T tests, anova tests, and linear mixed effect models were implemented in R.

### Bulk ATAC-seq processing

ATAC-seq data was generated from cryopreserved samples at 0h and 240h post IR. Cryopreservation medium was made using 90% FBS (R&D Systems, S11550H) and 10% DMSO (Fisher scientific, 31-761-00ML) solution. Healthy and IPF lung fibroblasts were seeded at 0.4 million to 1 million cells prior to IR, detached using Accutase (Sigma, A6964-500ML) at the collection timepoint, and sent to Novogene for QC prior to sequencing in cryopreservation medium. Samples were screened by Novogene for having greater than 50,000 cells and greater than or equal to 60% viable cells. All samples passed the screening criteria and were processed on the Novaseq (PE150) (Illumina) at 50 million paired reads per sample by Novogene.

FASTQ files were aligned using Bowtie2 against the hg38 reference genome^60,61^. Resulting SAM files were processed using samtools to produce sorted and deduplicated BAM files^62^. MACS3 tool was used to perform peak calling on the deduplicated BAM file^63^.

Bedtools was used on the resulting narrowPeak files resulting from peak calling to merge overlapping peaks across all samples to produce a single consensus set of peak regions^64^.

For each sample, a peak count matrix was created using the featureCounts tool from the SubRead toolkit to count the occurrences of reads per sample that intersect peaks^65^. Resulting peak counts were assembled into an AnnData object for further analysis using Differential Expression and GSEA methods outlined below (Supp. Fig. 5).

### Differential Expression Analysis

Differential expression analysis was performed using custom Python pipelines implementing a DESeq2-equivalent negative binomial generalized linear model framework (PyDESeq2), with a selective combination of disease groups (IPF, Healthy), treatment groups (IR, CTL) and time-points (0h, 8h, 24h, 48h, 240h) used as the primary variables in the design matrix for each contrast^66^. The analysis followed the canonical DESeq2 pipeline and subsequent estimation of coefficients and log2 fold-changes via Wald tests^67,68^. Genes were flagged differentially expressed at Benjamini–Hochberg false discovery rate (FDR) < 0.05 and |log2 fold-change| ≥ 0.4. Overlaps between differentially expressed genes (DEGs) across contrasts were visualized using UpSet plots generated with the pyUpSet package^69^.

Pathway-level changes were assessed by gene set enrichment analysis (GSEA) applied to the differential expression results, using gene sets from the Molecular Signatures Database (MSigDB) hallmark collection and the WikiPathways resource^70,71^. Full pathway enrichment across all time points can be found in Supp. Fig. 8.

### Fully Automated Senescence Test

FAST was performed as previously described with a detailed protocol here: https://www.protocols.io/view/fully-automated-senescence-test-fast-kxygx3ypwg8j/v1^30^. No positive control was used but negative controls without EdU incubation and minus SA-β-gal substrate were included in each FAST run.

### Immunostaining

Cells were fixed with 4% paraformaldehyde for 15 minutes, and then washed three times with PBS followed by permeabilization and blocking with 10% normal goat serum (Life Technologies, 50062Z) in 0.1% Triton X-100 (Sigma-Aldrich, X100RS-5G) for one hour. Cells were then incubated with primary antibodies at 4°C overnight (γH2AX, abcam, ab81299, 1:250; RAD51, abcam, ab133534, 1:1000; pATM, Life Technologies, MA1-2020, 1:250; PCNA, abcam, ab29, 1:400). The next day, after washing three times with PBS for 5 minutes each wash, secondary antibodies were applied for 1 hour. Cells were then washed once with PBS, counterstained with DAPI (Sigma, D9542-1MG) at 0.5 μg/ml in MQ water for 15 minutes, and washed once with MQ water. Finally, the cell solution was replaced with PBS before imaging. All steps performed at RT except for primary antibody incubation overnight.

### Image acquisition and analysis

Cells were plated on glass-based 96-well plates (VWR, MSPP-P9615HN). Images were taken on Harmony Operetta CLS type HH1600. Images were taken at 63X with water immersion, NA 1.15, and two peak autofocus. Confocal imaging was not used. Camera ROI was set to 2160×2160. Binning was set to 2. Images were captured using the following channels: DAPI, Alexa 488, and Alexa 647. For each cell and staining combination, 3 wells with 3 fields of view per well were captured. Each image is composed of 5 stacks with 0.5μm between stacks for a total height of 2μm. The average number of nuclei quantified per well was 27. Additional images from Fig. 3 can be found in Supp. Fig. 9.

Images were analyzed using Harmony PhenoLOGIC 5.2.2180.466. Images were analyzed with Basic Flatfield Correction with stacks processed using Maximum Projection. Nuclei were identified with the DAPI channel using Harmony “Find Nuclei” set to “Method B” with common threshold > 0.40 and area > 30μm². Background was subtracted for each channel with the formula A-quantile(A, 0.10).quantile, with negative values set to zero and undefined values set to “Local Average”. Intensity properties were calculated with the “Calculate intensity properties” building block. Intensity was quantified within the nuclear region and the mean fluorescence intensity for each well was normalized to the number of nuclei quantified. Spots were defined using “Find Spots” using “Method A”, specifically identifying spots within the nuclear region and normalized to the number of nuclei quantified in each well, with relative spot intensity > 0.040 and splitting sensitivity = 1.000.

### Weighted Gene Co-expression Network Analysis

Weighted Gene Co-expression Network Analysis was run using pyWGCNA^72^. pyWGCNA was run against the gene expression matrix in transcripts per million (TPM) format using the combination of irradiation treatment timepoints (0h, 8h, 24h, 48h, 240h) and sample source (IPF, HEALTHY) as sample groupings. Resulting gene modules were tested for gene set enrichment analysis (GSEA) against GOTerms (Gene Ontology), and cross referenced with differentially expressed genes (DEG). Sufficient power was reached using 16 total donors across multiple timepoints with two biological replicates per experiment.

### Single-cell Reference Data

Single-cell RNA-seq reference data from lungs with interstitial lung disease (ILD), including idiopathic pulmonary fibrosis (IPF), was obtained from Habermann et al. (GEO: GSE135893)^25^. Count matrices and accompanying metadata files were retrieved computationally from GEO and used with only minimal preprocessing required for integration with our analyses. Raw counts were normalized to counts per 10,000 (CP10k), and clustered using graph-based methods (louvain and leiden) to define transcriptomically distinct cell populations and clusters^73,74^. The mesenchymal compartment was highlighted by examining the gene set scores of IR-associated differentially expressed genes (IR DEGs) across these clusters.

### Geneset Scoring

A Seurat-style gene set scoring algorithm implemented in Scanpy was used to quantify enrichment of irradiation-associated differentially expressed genes (IR DEGs) within single-cell clusters^75,76^. For each cluster, a continuous score was derived from normalized expression values to summarize the relative expression of the IR DEG set. Higher scores indicate stronger similarity between the cluster-level expression profile and the IR-associated gene expression pattern. The IR gene signature was developed from the top 50 genes by fold change in the senescent healthy and IPF data sets which had consensus with single-cell mesenchymal interstitial lung disease (ILD) versus control differential expression. Bubbles generated above the heatmap show expression of each gene from the scRNA-seq data based on the IR gene signature. Boxplots provided in Supp. Fig. 3 shows the enrichment of the IR gene signature in the scRNA-seq data based on disease stratification. The IR gene signature consists of 163 genes, a subset of the entire time course healthy IR and IPF IR DEG list, and were identified as genes that had the most consensus signature to ILD from the scRNA-seq data. These 163 genes were compared to the SASP WikiPathway #3391 and SAUL_SEN_MAYO M45803 gene set using the ggvenn package (https://github.com/yanlinlin82/ggvenn) in R.

### Code availability

All analysis code, figure-generation notebooks and workflow utilities used in this study are available in the GitHub repository https://github.com/rubedolife/rb-buck-ipf-sen-2025. The repository will be made publicly accessible at the time of manuscript publication. rnaseqmut script will also be made available in a GitHub repository upon acceptance of the manuscript.

## DATA AVAILABILITY

Raw sequencing data (FASTQ files) and processed bulk RNA-seq and ATAC-seq data generated in this study will be deposited in the Gene Expression Omnibus (GEO); accession numbers will be provided upon acceptance of the manuscript. Combined metadata will also be made available upon final manuscript submission. All other data used in this study were obtained from previously published, publicly accessible resources, with source identifiers specified in the manuscript.

## Supporting information

20261126_Hughes_IPFpaper_Supplemental

Supplementary Data Donor Information

## ACKNOWLEDGEMENTS

We would like to acknowledge Dr. Judith Campisi and her legacy she built in the field of senescence that we hope to continue through this paper. We would like to thank the Buck Institute for letting the lab continue until we finish publishing papers started by Judy. This work was partially supported by P01AG066591 P-1, Training Grant - NIA T32 AG052374, and Rubedo Life Sciences. All schematics/diagrams were made using BioRender.

## AUTHOR CONTRIBUTIONS

JWB contributed to all parts of the experiments, analysis, and manuscript. AP contributed to all of the bioinformatics pipelines, developed figures, and helped write the methods. AS performed cell culture, ICC, and analysis. DC and LBM performed cell culture and RNA collection. AP and IJ helped with ATAC-seq analysis and the WGCNA. RB helped with cell culture and RNA collection. CH helped with the RNA-seq analysis. KS and FS helped with the rnaseqmut script. IB provided support on the CLS HTS. ALC provided cells from Cedars-Sinai. TCH provided cells from UCSF. HD helped with RNA collection. CJLS provided cells from UCSF and advised on the project. SDE, LME, and AV advised on the project. DF runs the bioinformatics core at the Buck where KS and FS performed analyses. CMH oversaw the provision of cells from Cedars-Sinai and edited the manuscript. AL contributed to and led all bioinformatics analyses. PYD and MQ oversaw the project and edited the manuscript. MQ and JC initiated the collaboration with the Buck Institute and Rubedo Life Sciences and JC oversaw the project until her passing in January 2024. All authors reviewed the manuscript.

## COMPETING INTERESTS POLICY

Authors affiliated with Rubedo Life Sciences are employees and/or hold equity or other financial interests in the company. These interests may be perceived to influence the work reported in this manuscript.

