## Supplementary material for "Uncovering senescent fibroblast heterogeneity connects DNA damage response to idiopathic pulmonary fibrosis": 20261126_Hughes_IPFpaper_Supplemental

Supplemental Figure 1. Fully-Automated Senescence Test (FAST) validates model of senescence induction by IR.

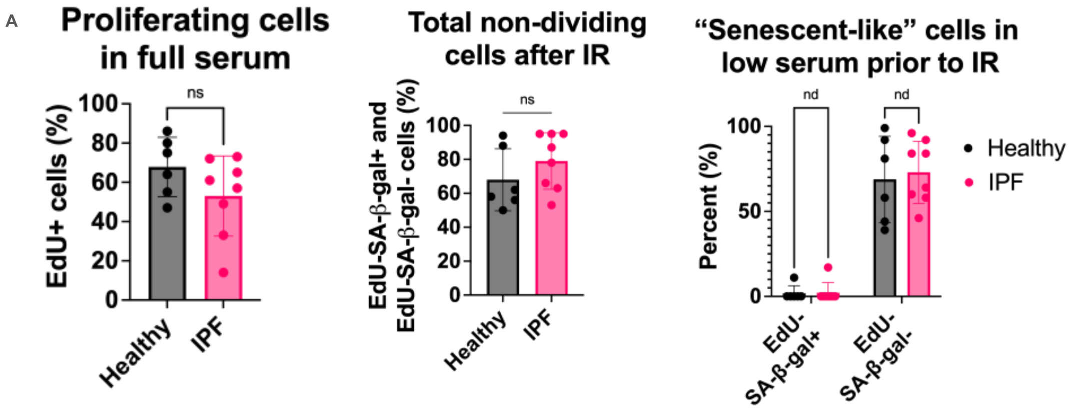


(A) Left panel shows healthy and IPF lung fibroblasts divide at similar rates. Middle panel shows that IR induces a largely homogeneous senescent-like phenotype by quantifying both EdU- (5-ethynyl-2′-deoxyuridine) SA-β-gal+ (senescence-associated beta-galactosidase) and EdU- SA-β-gal- populations. Right panel shows that prior to IR in low serum classically defined senescent cells (EdU-SA-β-gal+) are not present. 6 healthy and 8 IPF donors quantified for each plot (donors HLF872 and 99B omitted as these donors were not imaged and analyzed). 4 technical replicates of each condition was performed and averaged per donor. Unpaired and multiple unpaired t tests used with p value and q value cutoffs, respectively, set at < 0.05.

Supplemental Figure 2. Single nucleotide polymorphism (SNP) analysis shows little differences between healthy and IPF lung fibroblasts after IR.

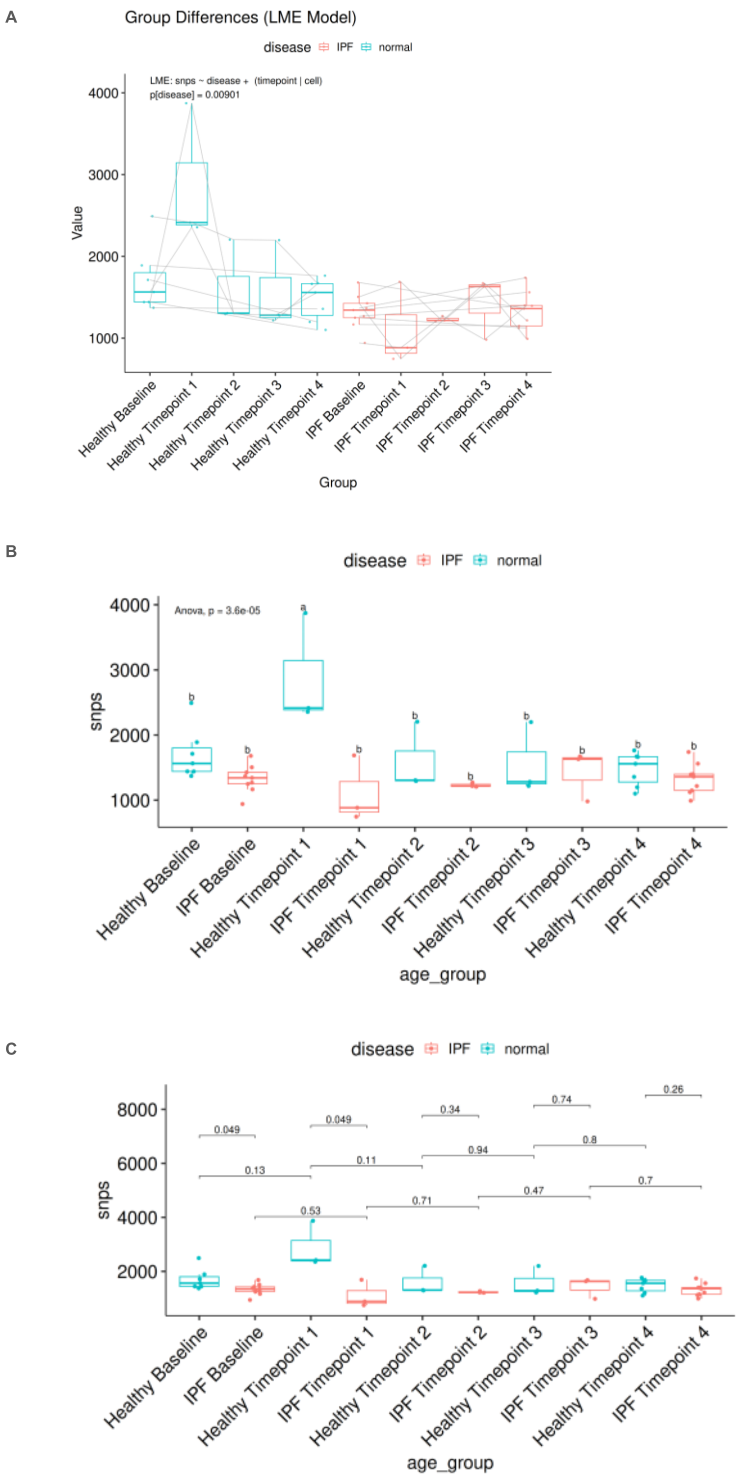


(A/B/C) RNA seq mutations across timepoints and disease classification identified using the package rnaseqmut and compared in R using a linear mixed effect model (A), anova (B), and t-tests (C).

Supplemental Figure 3. Scoring algorithm enrichment with disease stratification from healthy IR and IPF IR conditions.

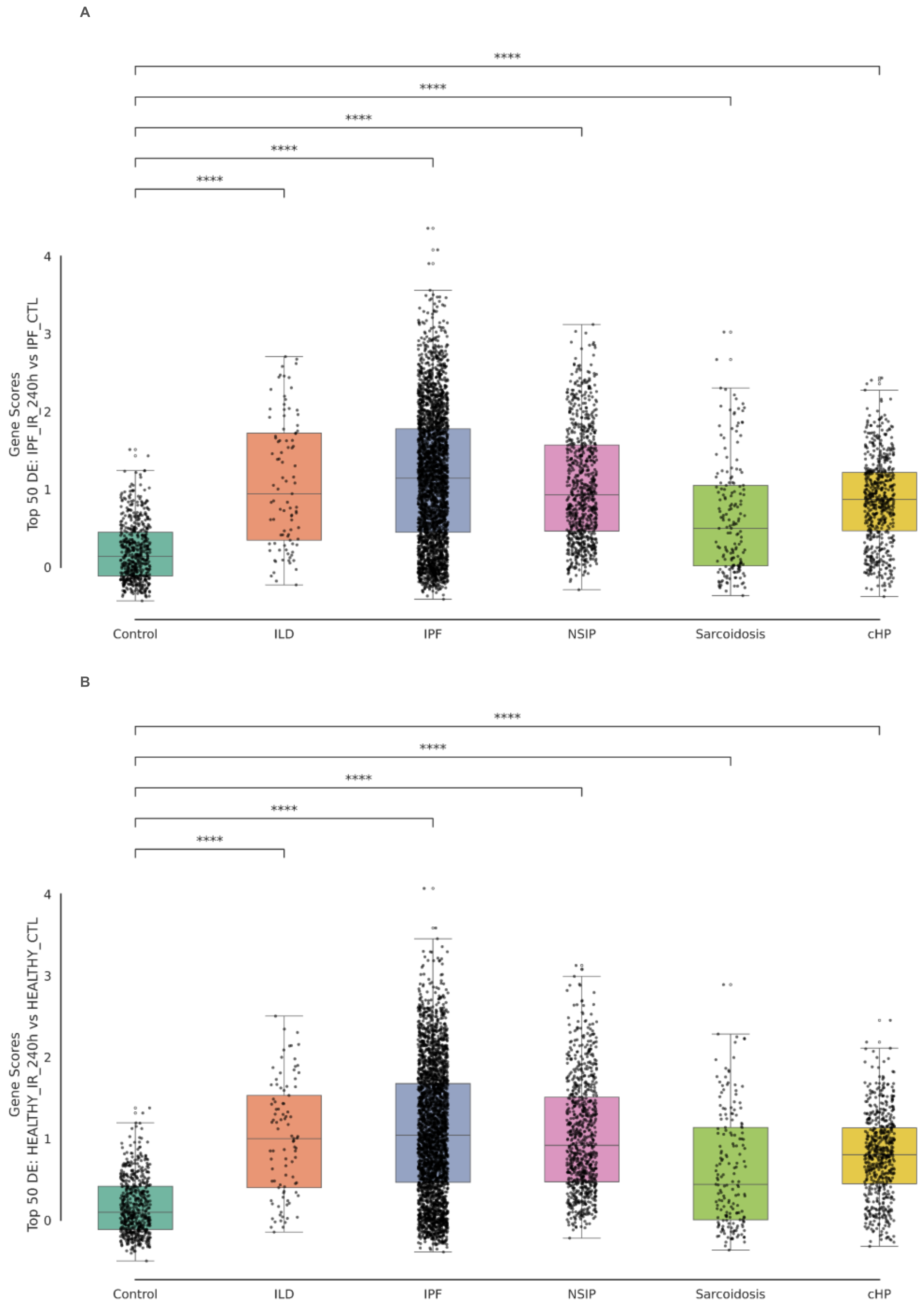


(A-B) Enrichment of our top 50 differentially expressed genes in our IPF IR (A) and healthy IR (B) signature against diseases stratified from the Habermann et. al. 2020 scRNA-seq study (IPF: Idiopathic Pulmonary Fibrosis, NSIP: Nonspecific Interstitial Pneumonia, Unclassifiable ILD: interstitial lung disease, cHP: Chronic hypersensitivity pneumonitis).

Supplemental Figure 4. MDM2 inhibition in IPF lung fibroblasts after IR shows a decrease in damage sites and intensity but also in protein repair intensity and cell counts.

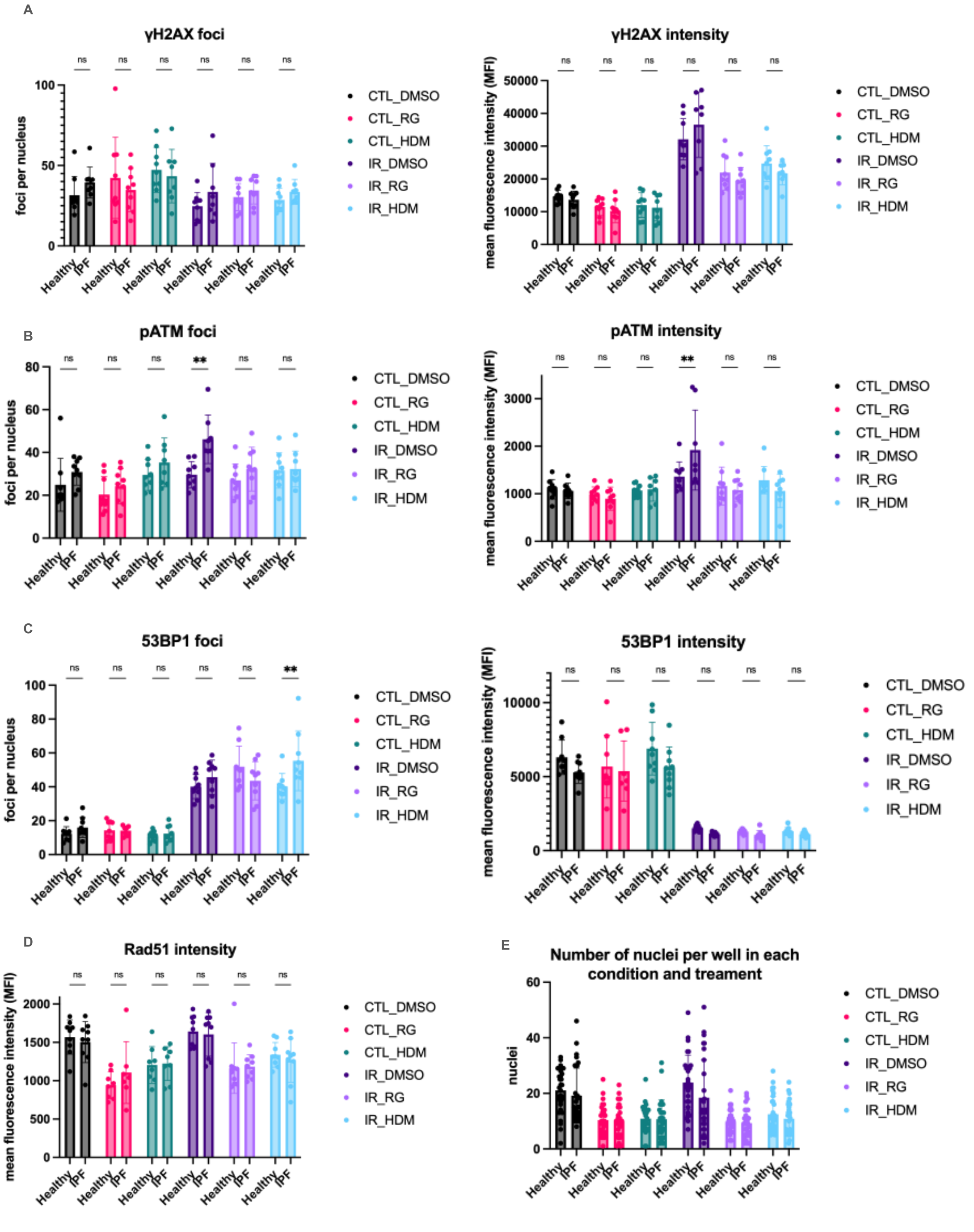


(A-E) 100nM of RG7388 or 25nM of HDM201 was used to treat healthy (donors: HLF250, HLF763, HLF872) or IPF (donors: 83A, 112A, 155B) lung fibroblasts 5 days prior to IR and cells were fixed for ICC 2 hours after IR. 3 technical replicates of each condition was performed. γH2AX (A), pATM (B), and 53BP1 (C) foci and intensity were quantified using the Harmony imaging and analysis method. Rad51 (D) intensity and the number of nuclei were quantified using the Harmony imaging and analysis method as well. Statistics (A-D) were calculated using a two-way ANOVA comparing column means within each cell of the column which included interaction terms and corrected for multiple comparisons using the Šídák multiple comparisons test with alpha = 0.05.

Supplemental Figure 5. ATAC-seq of healthy and IPF lung fibroblasts after IR shows heterogeneity of senescence response.

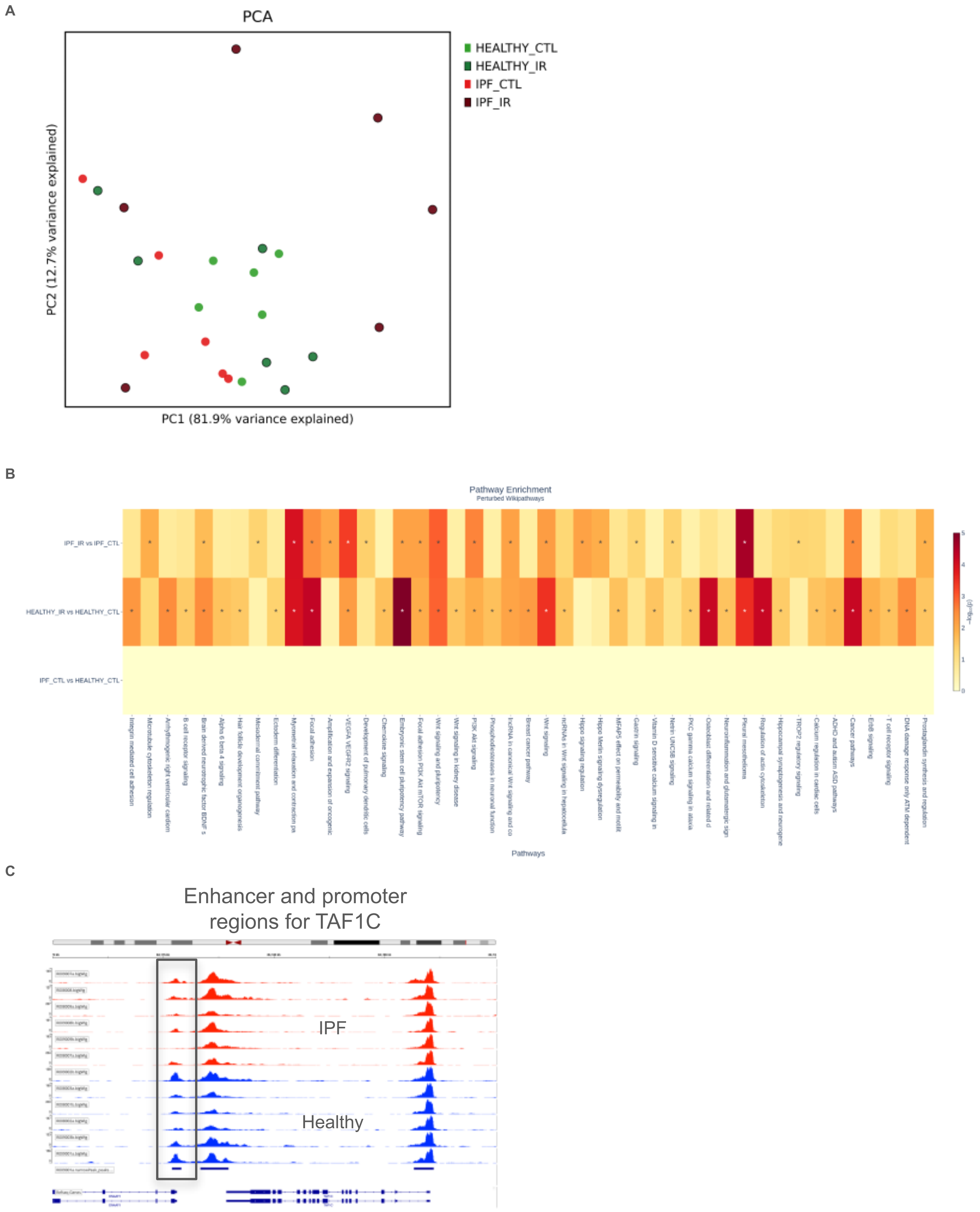


(A) PCA of healthy (donors: AJIQ, AJAZ AJDL, HLF250, HLF763, HLF872) and IPF (donors: 83A, 86A, 110A, 127A, 155B, IPF809) lung fibroblasts after IR does not seem to cluster in a specific pattern. (B) Pathway analysis of differentially accessible regions (DAR) in healthy and IPF lung fibroblasts with and without IR. DAR cutoffs were defined by log2FC < -1 or >1 and p-adjusted value < 0.05. (C) Chromatin tracks of healthy and IPF lung fibroblasts after 10 days after IR highlights TAF1C, a gene known to play a role in DDR and DNA repair after DNA damage. Box highlights an enhancer region in TAF1C lowered in IPF relative to healthy.

Supplemental Figure 6. ABT263 dose response curves in young, adult, and IPF lung fibroblasts.

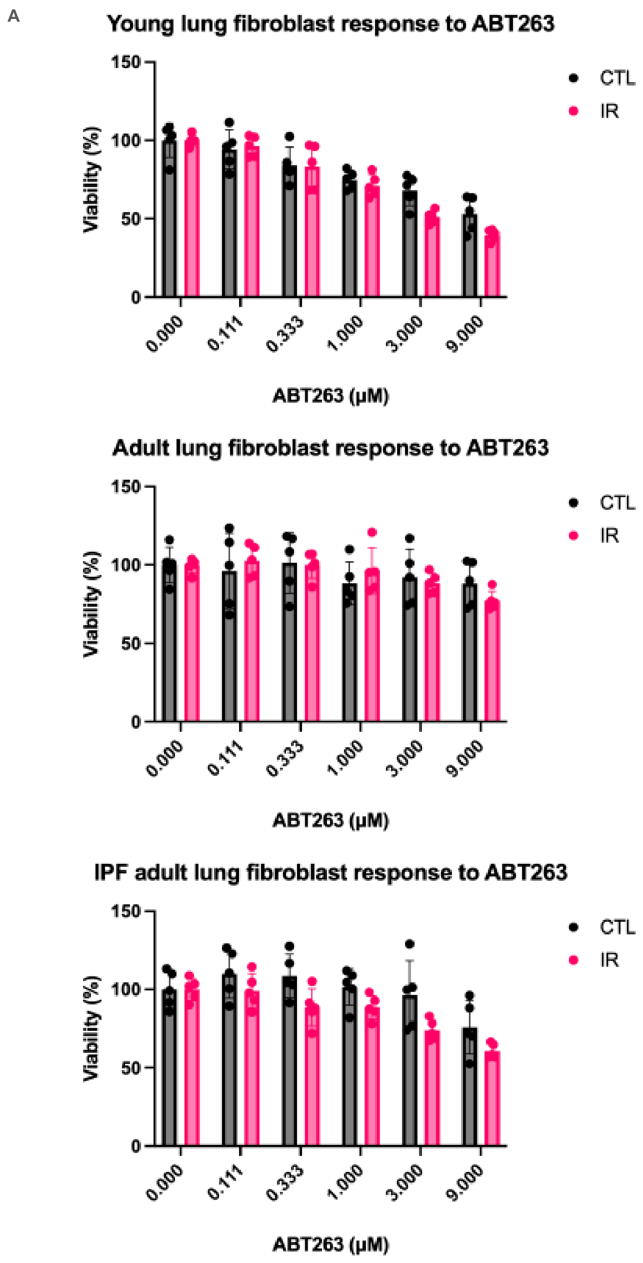


(A) ABT263 dose response performed 10 days after IR for 24 hours and assessed using the FAST protocol. DAPI staining was used to assess viability based on the CTL (0μM) DAPI cell counts. One donor (Young: IMR90, Adult: HLF763, IPF: 86A) was used with 4 technical replicates per concentration.

Supplemental Figure 7. Schematic of experimental design.

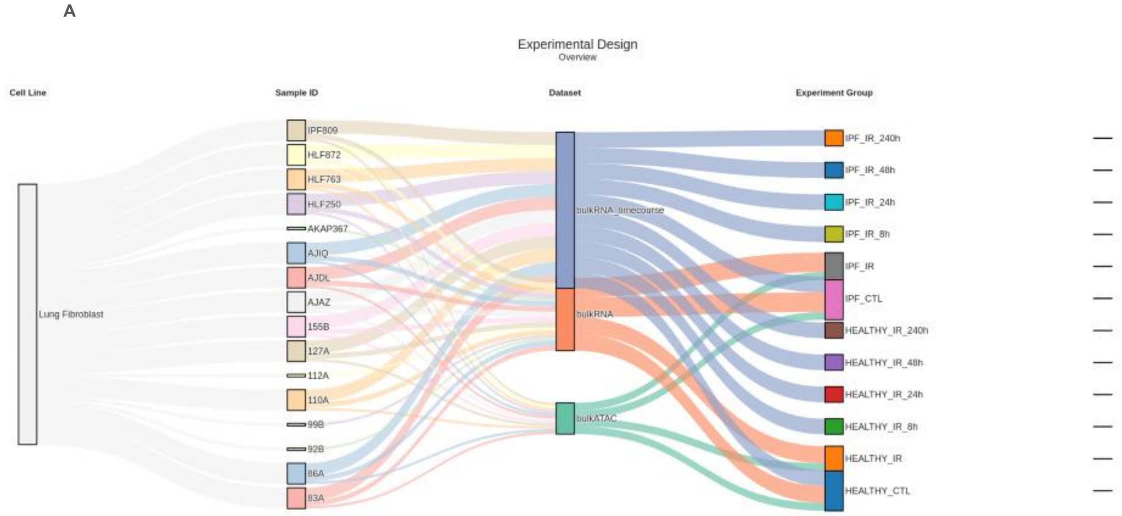


(A) Cell line is primary, human lung fibroblasts. Sample ID refers to the donor ID label which can be found in the “Supplementary Data Donor Information” table. All samples were in full serum (FS) for RNA seq and ATAC-seq. 6 healthy and 6 IPF donors were used for time course RNA seq and ATAC-seq (Sample IDs: AJAZ, HLF250, AJDL, HLF872, HLF763, AJIQ, 83A, 110A, IPF809, 86A, 127A, and 155B). All sample IDs processed 10 days after IR.

Supplemental Figure 8. RNA sequencing after IR over time shows heterogeneous responses to senescence induction within the same cell type dependent on disease.

A

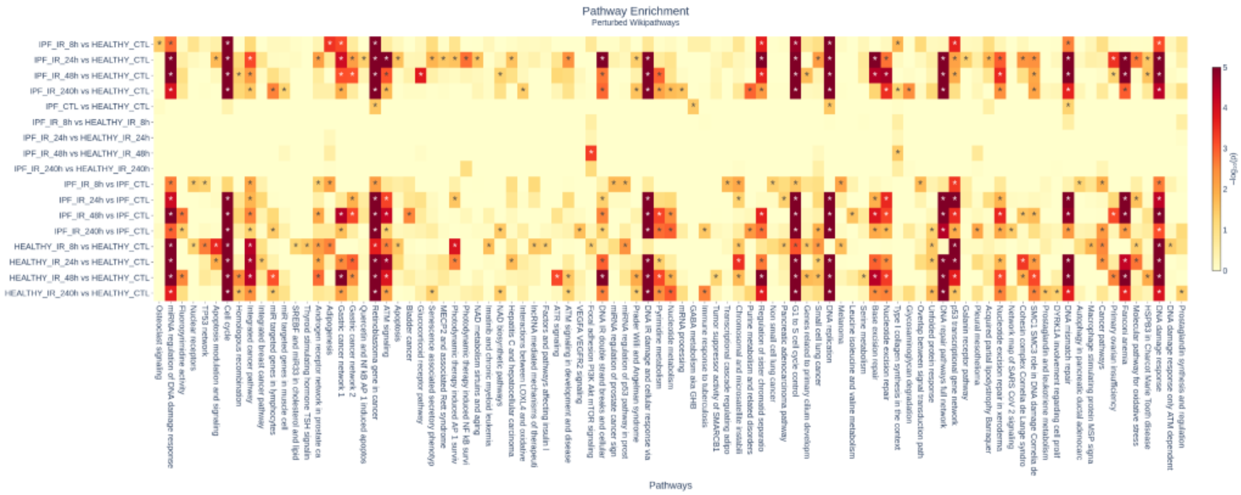


(A) Gene set enrichment analysis performed using Wikipathways of DEGs in healthy and IPF lung fibroblasts over time after IR.

Supplemental Figure 9. pATM and PCNA staining of healthy and IPF lung fibroblasts prior to and over time after IR.

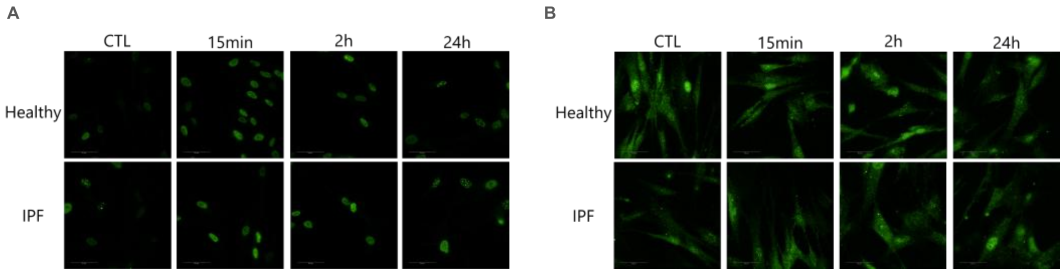


(A) pATM imaging quantified in Fig. 3E-F. (B) PCNA imaging quantified in Fig. 3E-F.
